# A high-resolution mass spectrometry-based method for quantifying insulin-stimulated glucose uptake in mice following an intraperitoneal injection of tracer

**DOI:** 10.64898/2026.03.31.714892

**Authors:** Guo-Fang Zhang, Dorothy H. Slentz, Louise Lantier, Owen P. McGuinness, Deborah M. Muoio, Ashley S. Williams

## Abstract

**Objective:** A catheter-free, non-radiolabeled method that permits *in vivo* measurement of tissue-specific glucose uptake does not exist. To address this gap, we sought to develop and validate a new, higher throughput mass spectrometry (MS)-based method that combines an injection of insulin with a non-radiolabeled glucose tracer, 2-fluoro-2-deoxyglucose (2FDG), to determine insulin-stimulated tissue-specific glucose clearance in conscious, unrestrained mice.

**Methods:** Injections of saline or insulin with 2FDG were coupled with LC-Q Exactive Hybrid Quadrupole-Orbitrap (LC) MS-based measures of plasma 2FDG and tissue (2-fluoro-2-deoxyglucose-6-phosphate) 2FDGP to determine glucose clearance in mice under several different conditions.

**Results:** The newly developed method was first applied to a dose response experiment in mice. Next, the ability of this method to quantify changes in glucose clearance in response to an insulin stimulus was assessed, and glucose clearance was compared between chow and high fat fed mice. Results from these studies showed that insulin-stimulated skeletal muscle and heart glucose clearance can be estimated following a bolus injection of tracer, and these fluxes are blunted in diet-induced obese mice. The broad applicability of this approach was then demonstrated by assessing glucose clearance in a mouse model with anticipated changes in insulin-stimulated skeletal muscle glucose metabolism.

**Conclusions:** The results validated a new LC-MS method to quantify insulin-stimulated tissue-specific glucose clearance *in vivo* without the use of catheters or radiolabeled tracers. The method offers great potential because it is designed for application to pre-clinical studies seeking high throughput tests and/or assays that can be coupled with discovery technologies such as genomics, proteomics and metabolomics.

**HIGHLIGHTS:**

- *In vivo* glucose clearance can be estimated by a new non-radiolabeled method.
- The plasma tracer to tracee ratio is required to determine tissue tracer phosphorylation.
- Measures of plasma glucose and tracer kinetics are critical for data interpretation.
- The new method can be combined with ‘omics’ technologies such as metabolomics.

## 1. INTRODUCTION

Advancements in metabolic research are made possible by mouse models of human disease and the miniaturization of clinical tests. Among these tests, one of the most established is the hyperinsulinemic-euglycemic (insulin) clamp, which uses labeled glucose analogues in conscious, catheterized, unrestrained mice to determine sites of insulin resistance [1, 2]. Within the insulin clamp, peripheral tissue insulin resistance is assessed by combining radiolabeled tracers, such as 2-[^14^C(U)]deoxy-D-glucose, with the in vivo 2DG method. This method involves a bolus injection of 2DG in trace amounts (e.g. 13 µCi radiolabeled 2DG or ∼237nmol) followed by serial collection of blood samples to measure blood glucose and rates of 2DG disappearance from the plasma [3–5]. The plasma 2DG decay curve and measures of plasma glucose are then coupled with measures of tissue 2-deoxyglucose-6-phosphate (2DGP) to calculate the tissue-specific clearance of 2DG (*K*_g_) or glucose metabolic index (*R*_g_) [4].

Whereas the insulin clamp is likely to maintain gold standard status for many years to come, this method has several caveats that limit its broad utility. Prominent among these is its heavy reliance on indwelling catheters and radiolabeled tracers. The foregoing caveats tend to push scientists towards other less technical approaches, such as glucose and insulin tolerance tests; however, these tests do not measure insulin action nor do they provide information regarding peripheral insulin action [2, 6, 7]. Accordingly, the field has long recognized a need for an ‘intermediate’ phenotyping test that can be performed alongside glucose or insulin tolerance tests to identify tissue-specific alterations in glucose or insulin dependent regulation. This has led several groups to employ a modified version of the radiolabeled *in vivo* 2DG method that omits the use of catheters [8–13]. However, most studies only report tissue radioactivity [8, 12] or tissue 2DGP levels [11]; and these are not quantitative measures of glucose uptake without the accompanying plasma glucose levels, plasma 2DG decay curve, and tissue 2DGP levels from which *K*_g_ and *R*_g_ can be calculated.

Nonetheless, the limitations of radiolabeled tracers can be overcome by the use of mass-spectrometry (MS)-based approaches that deploy non-radiolabeled glucose analogues. For this reason, we developed and validated a new LC-MS based method to measure trace amounts of the established glucose analog, 2-fluoro-2-deoxyglucose (2FDG) and the phosphorylated metabolite, 2-fluoro-2-deoxyglucose-6-phosphate (2FDGP) in biological samples [14]. In this study, we applied this method and further developed a new LC-MS based method to measure insulin-stimulated glucose uptake in unrestrained, conscious mice following a bolus injection of 2FDG with insulin. Our new mouse metabolic phenotyping test offers several advantages over conventional approaches as it is a higher throughput option for measuring *in vivo* tissue glucose metabolism and allows the use of biological specimens for secondary analyses, including state-of-the-art “omics” technologies.

## 2. METHODS

### 2.1. Chemicals and reagents

2DG was purchased from Chem-impex Int’l Inc (Wood Dale, IL). 2DGP and 2FDG were purchased from Santa Cruz Biotechnology (Dallas, Texas). 2FDGP was purchased from Omicron Biochemicals Inc. (South Bend, IN). 2FDG, 2-chloro-2-deoxyglucose (2ClDG), hexokinase from baker yeast, triethanolamine, formic acid, chloroform, adenosine triphosphate (ATP), ethylenediaminetetraacetic acid disodium salt (EDTA), and other chemical reagents in analytical grade or above were purchased from Sigma (St. Louis, MO). 2ClDGP was synthesized in house. Briefly, 150 ul of 100 mM 2ClDG, 180 ul of 100 mM ATP, 180 ul of 100 mM MgCl_2_, 75 ul of 100 mM EDTA, and 10 U hexokinase were added into 1 ml triethanolamine buffer (50 mM, pH 7.4). The reaction was maintained at 37°C for 30 min. The reaction was quenched by adding 1 ml of pre-cooled methanol. After centrifugation at 800 × g for 15 min, the supernatant was dried and dissolved in distilled water. The concentration of 2ClDGP was estimated based on 2DGP. 2ClDGP was used as the internal standard (IS) for the external calibration curve, therefore, the absolute concentration was not required.

### 2.2. Mouse models and experimental design

All animal studies were approved by the Duke University or Vanderbilt University Institutional Animal Care and Use Committee (IACUC) and conducted in Association for Assessment and Accreditation of Laboratory Animal Care-associated (AAALAC) facilities. Mice were housed in a light (12h light/12h dark) and temperature (22°C) controlled room and had *ad libitum* access to food and water unless noted otherwise. At the end of each study, mice were anesthetized with Nembutal (intraperitoneal (i.p.) injection; 100 mg/kg body weight) prior to organ removal. Female mice were excluded due to established sex differences in body weight and insulin action [15] and reviewed in [16].

#### 2.2.1. Hyperinsulinemic-euglycemic (insulin) clamp (IC) with radiolabeled (^14^C) 2DG

A comprehensive step-by-step description of the insulin clamp surgery, isotope clamp method, and calculations is available from the Vanderbilt Mouse Metabolic Phenotyping Center (MMPC) website (https://Vmmpc.org). Insulin clamp studies were performed in chronically catheterized, conscious, unstressed, 5h fasted mice at the Vanderbilt University MMPC. Carotid artery and jugular vein catheters were surgically implanted 5-7 days prior to study. Mice were weighed daily and if mice lost more than 10% of their pre-surgery body weight, they were excluded from the study. On the day of study, mice were fasted and [3-^3^H]glucose was primed and continuously infused into the jugular vein from t= −90 to t=0 min (0.04 μCi/min). Prior to the start of the insulin clamp, arterial blood samples were taken at t= −15 and t= −5 min for the determination of basal arterial blood glucose, insulin, and [3-^3^H]glucose turnover. The insulin clamp was started at t= 0 with a continuous insulin infusion (4mU/(kg•min)) and a variable infusion of glucose+[3-^3^H]glucose (0.06 μCi/μL) to maintain euglycemia. The insulin infusion rate was chosen to assess muscle insulin action [1]. A mixture of glucose+[3-^3^H]glucose was employed to minimize alterations in [3-^3^H]glucose specific activity upon changes in the glucose infusion rate (GIR). Heparinized, saline washed erythrocytes were infused to prevent a fall in hematocrit. Arterial glucose was monitored by a hand-held glucometer (AccuCheck) every 10 min and the GIR was adjusted to maintain and clamp arterial glucose. Arterial samples for the determination of clamp glucose and [3-^3^H]glucose turnover were collected at t= 80, 90, 100, and 120 min, and samples for the determination of clamp insulin were collected at t= 100 and 120 min. At t= 120 min, a 13μCi bolus of 2[^14^C]deoxyglucose ([^14^C]2DG) was administered into the venous line to determine tissue glucose uptake or the glucose metabolic index (*R*_g_). Blood samples were collected at t= 122, 125, 135, 145, and 155 min to determine [^14^C]2DG disappearance from the plasma. After the final blood sample was collected, mice were anesthetized and tissues were freeze clamped and stored at −80°C until analysis.

#### 2.2.2. Intraperitoneal (i.p.) injection of saline or insulin with or without unlabeled 2DG

Chow fed mice on a C57BL/6NJ background were habituated for 3 days by i.p. injections of saline. On the day of study, mice were fasted on Alpha-dri bedding for 5h with access to water and injected i.p. with either saline or insulin (1.5 U/kg body mass) with or without 25 µmol 2-deoxyglucose (2DG). Blood samples were obtained from the tail at baseline (t= 0 min) and at t= 10, 20 and 30 min after injection and used for measures of blood glucose via hand-held glucometer (Bayer Contour Blood Glucose Monitoring System).

#### 2.2.3. Hyperinsulinemic-euglycemic (insulin) clamp (IC) with unlabeled 2DG

Insulin clamp studies were performed in chronically catheterized, conscious, unstressed, 5h fasted male mice on a C57BL/6J background at the Vanderbilt University MMPC. Carotid artery and jugular vein catheters were surgically implanted 5-7 days prior to study. Mice were weighed daily and if mice lost more than 10% of their pre-surgery body weight, they were excluded from the study. Prior to the start of the insulin clamp, arterial blood samples were taken at t= −15 and t= −5 min for the determination of basal arterial blood glucose. The insulin clamp was started at t= 0 with a continuous insulin infusion (4mU/(kg•min)) and a variable infusion of glucose to maintain euglycemia. The insulin infusion rate was chosen to assess muscle insulin action [1]. Heparinized, saline washed erythrocytes were infused to prevent a fall in hematocrit. Arterial glucose was monitored by a hand-held glucometer (AccuCheck) every 10 min and the glucose infusion rate (GIR) was adjusted to maintain and clamp arterial glucose. At t= 120 min, a 20 µmol bolus of 2DG was administered into the venous line. Importantly, the GIR was not changed after 2DG administration. Blood samples were collected at t= 122, 125, 135, 145, and 155 min to determine blood glucose and plasma 2DG.

#### 2.2.4. 2FDG dose response

Chow fed mice on a C57BL/6NJ background from Jackson Laboratory (strain #005304) were habituated for 3 days by i.p. injections of saline. On the day of study, mice were fasted for 5 h with access to water and injected i.p. with saline+75 nmol 2FDG or saline+225 nmol 2FDG. Tail blood samples were obtained at t= 30 min after injection to determine plasma 2FDG, and mice were anesthetized and tissues were harvested and freeze clamped for measures of tissue 2FDGP.

#### 2.2.5. Hyperinsulinemic-euglycemic (insulin) clamp (IC) with unlabeled 2FDG

Insulin clamp studies were performed in chronically catheterized, conscious, unstressed, 5h fasted C57BL/6J male mice from Jackson Laboratory (strain #000664) at the Vanderbilt University MMPC. Carotid artery and jugular vein catheters were surgically implanted 5-7 days prior to study. Mice were weighed daily and if mice lost more than 10% of their pre-surgery body weight, they were excluded from the study. Prior to the start of the insulin clamp, arterial blood samples were taken at t= −15 and t= −5 min for the determination of basal arterial blood glucose. The insulin clamp was started at t= 0 min with a continuous insulin infusion (4mU/(kg•min)) and a variable infusion of glucose to maintain euglycemia. The insulin infusion rate was chosen to assess muscle insulin action [1]. Heparinized, saline washed erythrocytes were infused to prevent a fall in hematocrit. Arterial glucose was monitored by a hand-held glucometer (AccuCheck) every 10 min and the glucose infusion rate (GIR) was adjusted to maintain and clamp arterial glucose. At t= 120 min, a 225 nmol bolus of 2FDG was administered into the venous line. Importantly, the GIR was not changed after 2FDG administration. Blood samples were collected at t= 122, 125, 135, 145, and 155 min to determine blood glucose.

#### 2.2.6. Effect of route of 2FDG administration on glucose uptake

Male chow fed mice on a C57BL/6NJ background from Jackson Laboratory (strain #005304) were habituated for 3 days by i.p. injections of saline. On the day of study, mice were singly housed and fasted on Alpha-dri bedding for 5h with access to water. Mice were injected i.p. or i.v. (tail vein) with saline+225 nmol 2FDG or insulin (1.5U/kg body mass)+225 nmol 2FDG and tissue-specific glucose uptake was determined in the conscious, unrestrained state. Blood samples were obtained from the tail prior to and at t= 5, 10, 15, 25 and 35 min after injection and used for measures of blood glucose and disappearance of 2FDG from the plasma. Mice were anesthetized at t= 35 min and tissues were harvested, freeze clamped and stored at −80°C until subsequent analysis.

#### 2.2.7. Effect of diet on muscle glucose uptake

Studies were performed in chow and high fat (HF) fed C57BL/6J male mice from Jackson Laboratory (strains #000664 and #380050 respectively). Notably, HF fed mice were placed on a 60% kcal as fat diet (Research Diets D12492) starting at 6 weeks of age. At approximately 16 weeks of age, mice were weighed, and body composition and oral glucose tolerance were determined (see sections 2.4 and 2.5 below for information regarding body composition and oral glucose tolerance tests). For the assessment of tissue-specific glucose uptake, 17-week-old mice were habituated for 3 days by i.p. injections of saline. On the day of study, mice were singly housed and fasted on Alpha-dri bedding for 5 h with access to water prior to an i.p. injection with insulin (1.5U/kg lean mass) + 225 nmol 2FDG. Blood samples were obtained from the tail at t= 0 and t= 5, 10, 15, 25 and 35 min after injection and used for measures of blood glucose and disappearance of 2FDG from the plasma. Mice were anesthetized at t= 35 min and tissues were harvested, freeze clamped and stored at −80°C until subsequent analysis.

#### 2.2.8. Effect of skeletal muscle-specific deletion of thioredoxin-interacting protein (TXNIP) on muscle glucose uptake

TXNIP^fl/fl^ (floxed littermate controls or FC) and mice with a skm-specific deletion of TXNIP (TXNIP^Myo^ or mKO) mice on a C57BL/6NJ background were generated as described in {DeBalsi, 2014 #15. At approximately 13-16 weeks of age, mice were weighed, and body composition and oral glucose tolerance were determined (see sections 2.4 and 2.5 below for information regarding body composition and oral glucose tolerance tests). For the assessment of tissue-specific glucose uptake, approximately 20-week-old mice were habituated for 3 days by i.p. injections of saline. On the day of study, mice were singly housed and fasted on Alpha-dri bedding for 5 h with access to water prior to an i.p. injection with insulin (0.75U/kg body mass) + 225 nmol 2FDG. Blood samples were obtained from the tail before injection at t= 0 and at t= 5, 10, 15, 25 and 35 min after injection and used for measures of blood glucose and disappearance of 2FDG from the plasma. Mice were anesthetized at t= 35 min and tissues were harvested, freeze clamped and stored at −80°C until subsequent analysis.

### 2.3. Blood and plasma metabolites

Blood glucose was determined as previously described {Williams, 2022 #16}. Blood beta-hydroxybutyrate (3OHB) was determined using a Keto-Mojo Ketone Meter (Keto-Mojo). Blood lactate was determined using a Lactate Plus Meter (Nova Biomedical). Tail blood was collected using heparin or EDTA-coated capillary tubes (Sarstedt Inc. MICROVETTE CB300, Sarstedt Inc.) and plasma was isolated for the determination of circulating 2FDG.

### 2.4. Body composition

Body weight and body composition were measured in fed mice. Body composition was determined using a mq10 nuclear magnetic resonance analyzer (Bruker Optics).

### 2.5. Oral glucose tolerance tests

Mice were fasted on Alpha-dri bedding for 5h with access to water. At the start of the test, mice were weighed and a baseline blood sample was obtained from the tail for the determination of fasting blood glucose (Bayer Contour Blood Glucose Monitoring System). At t= 0 min mice received an oral gavage of a 2g/kg lean mass glucose solution (45% glucose diluted in water) for studies described in section 2.2.7 or 2g/kg body weight for studies described in section 2.2.8. Subsequent blood glucose samples were obtained via the tail at t= 15, 30, 60, and 90min post-gavage.

### 2.6. LC-Q Exactive Hybrid Quadrupole-Orbitrap Mass Spectrometer sample preparation

Plasma 2FDG and tissue 2FDGP were extracted using a dual-phase folch extraction process as described below.

#### 2.6.1. Plasma 2FDG analysis

Ten microliters (μl) of a 0.025mM 2ClDG internal standard solution was pipetted into a clean 2 ml tube. Plasma samples were thawed on ice and 2 μl of each sample was added prior to folch extraction with the following solvents: 200 μl ice cold methanol, 200 μl chloroform, and 200 μl distilled water with vortexing after each addition. The sample mixture was then centrifuged for 20 min at 14,000 ×g at 4°C and 350 μl of the upper phase was aliquoted into a clean 1.7 ml tube and dried completely by nitrogen gas at 32°C. The dried residue was resuspended in 30 µl distilled water, vortexed, and placed in an autosampler vial for LC-MS analysis.

#### 2.6.2. Tissue 2FDGP analysis

Tissues were pulverized under liquid nitrogen. Approximately 10 mg powdered tissue was weighed prior to folch extraction with the following solvents: 400 μl ice cold methanol + 10 μl of 2ClDGP internal standard (0.05 mM), 400 μl chloroform, and 400 μl distilled water using a Qiagen Tissuelyzer (30 Hz, 1 min per solvent). The internal standard was added after the methanol and prior to the chloroform and water to ensure sample loss was accounted for during the extraction. After homogenization, the sample mixture was centrifuged for 20 min at 14,000 ×g at 4°C. The upper phase (∼750 µl) was removed and used to generate two 350 μl aliquots. One aliquot was stored at −80°C and the second was dried completely by nitrogen gas at 32°C. The dried residue was resuspended in 100 µl distilled water, vortexed, and placed in an autosampler vial for LC-MS analysis.

### 2.7. LC-Q Exactive Hybrid Quadrupole-Orbitrap Mass Spectrometer instrumentation and conditions

The Thermo Scientific Vanquish UHPLC system was configured with a binary pump, a thermostatted column compartment, and a temperature-controlled autosampler. The binary pump was used to transport mobile phase (HPLC-MS grade water) at a flow rate of 0.5 ml/min in isocratic elution mode. The column was a Microsorb-MV C18 column (100 × 4.6 mm, 3 µm) with C18 guard column and was kept at 40°C in the oven compartment. The autosampler was maintained at 5°C, and the injection volume was 1 µl. The total running time was 7 min. The parameters for Q-Exactive^+^-MS equipped with a HESI probe: heat temperature: 425°C; sheath gas: 30, auxiliary gas, 13; sweep gas, 3; spray voltage, 3.5 kV for positive mode; capillary temperature was set at 320 °C, and S-lens was 45. A full scan range was set at 60 to 900 (m/z). The resolution was set at 70,000 (at m/z 200). The maximum injection time (max IT) was 200 ms. Automated gain control (AGC) was targeted at 3 × 10^6^ ions.

### 2.8. LC-Q Exactive data analysis

Mass spectrometric data was acquired by Thermo Xcalibur software. Analyte concentrations were calculated using a 1/x weighted linear regression analysis of a standard curve [14].

### 2.9. Calculation of the tissue-specific clearance of 2FDG (*K*_g_) and tissue-specific glucose metabolic index (*R*_g_)

*K*_g_ (mL/min/100g tissue) and *R*_g_ (µmol/min/100g tissue) were calculated from measured tissue 2FDGP (µmol/mg), plasma 2FDG area under the curve (AUC 2FDG plasma; µM•min), and average blood glucose (μmol/ml) during the period after 2FDG injection. The equations for *K*_g_ and *R*_g_ are as described in [4] and as follows:

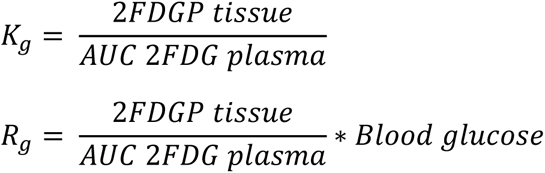

Importantly, the t=0 min timepoint was excluded for measures of blood glucose when calculating *R*_g_. *R*_g_’ was calculated as follows:

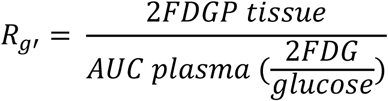

### 2.10. Gene expression

RNA was isolated from approximately 20 mgs of frozen, pulverized gastrocnemius (Gastroc), quadriceps (Quad), heart and liver tissues using a Trizol/chloroform extraction method paired with RNeasy Mini spin columns (Qiagen) with on column DNase treatment. RNA concentration was determined using a NanoDrop (Thermo Scientific). DNA was synthesized using the iScript cDNA Synthesis Kit (Bio-Rad). Quantitative PCR (qPCR) was performed using IDT Primetime Master Mix and the following assays: Txnip (Mm01265659_g1) and 18S (Applied Biosystems 4318839). mRNA quantities were determined by comparing the reported Ct value of each target/reference gene to a standard curve performed in the same RT-PCR run. Data were normalized to 18S.

### 2.11. Targeted mass spectrometry-based metabolomics

Tissue and plasma acylcarnitines and amino acids were determined as previously described [17].

### 2.12. Statistical analysis

Data are presented as means ± SEM. Statistical analyses were performed using GraphPad Prism 9.0 (GraphPad Software, San Diego, CA) using two-tailed student’s t-tests. Figures were generated using GraphPad Prism 9.0. The level of significance for all experiments was set at P ≤ 0.05.

## 3. RESULTS

### 3.1. Measurement of *in vivo* glucose uptake requires nanomolar quantities of glucose analogs

The goal of this study was to develop and validate a new, MS-based mouse phenotyping test to assess *in vivo* insulin-stimulated glucose uptake following a bolus injection of tracer. This new phenotyping test features a modified version of the 2DG method traditionally applied towards the end of a hyperinsulinemic-euglycemic (insulin) clamp (Figures 1A and 1B). The insulin clamp experimental setup and protocol are depicted in Figure 1B and described in detail in the Methods section 2.2.1. In the insulin clamp, indwelling arterial and venous catheters are inserted into mice; the arterial line is used for sampling and the venous line is used for infusions. Insulin is infused at a constant rate and unlabeled glucose mixed with radiolabeled glucose is infused at a constant rate to maintain euglycemia (Figure 1D). The amount of glucose infused (also known as the glucose infusion rate or GIR) reflects the insulin sensitivity of the mouse (Figure 1C). At the end of the insulin clamp, a 13 µCi bolus of ^14^C 2DG is administered in the venous line and blood samples are taken for the assessment of blood glucose and disappearance of 2DG. Once the study is completed, mice are anesthetized and tissues are collected for the determination of tissue ^14^C 2DGP. A critical, but often overlooked, consideration for these assays is that high (i.e. non-tracer) doses of 2DG can perturb whole body glucose metabolism. To this end, we first sought to determine if a bolus injection of 25 µmol unlabeled 2DG altered blood glucose levels in chow fed, unrestrained, conscious mice (Figure 1E). Our data showed that there was no difference in blood glucose between mice injected with saline or saline + 25 μmol 2DG (Figure 1F); and as anticipated, insulin precipitated a fall in blood glucose and this was not altered by the addition of 25 μmol 2DG (Figure 1F). Next, we wanted to determine if bolus administration of 20 μmol unlabeled 2DG altered blood glucose during an insulin clamp. Accordingly, we conducted insulin clamp experiments and administered 20 μmol 2DG in the venous line, in place of 13 µCi ^14^C 2DG. Importantly, for this experiment, the GIR was not adjusted after 2DG injection so the change in blood glucose reflected the impact of 2DG on systemic glucose homeostasis under insulin clamped conditions (Figures 1G-1H). From this experiment, we found that intravenous injection of 20 μmol unlabeled 2DG during an insulin clamp dramatically increased blood glucose (Figure 1I), presumably due to inhibition of hexokinase in the brain and a concomitant central counterregulatory response, that in turn impaired whole body glucose disposal (Figure 1J and referenced in [18]).

**Figure 1.**
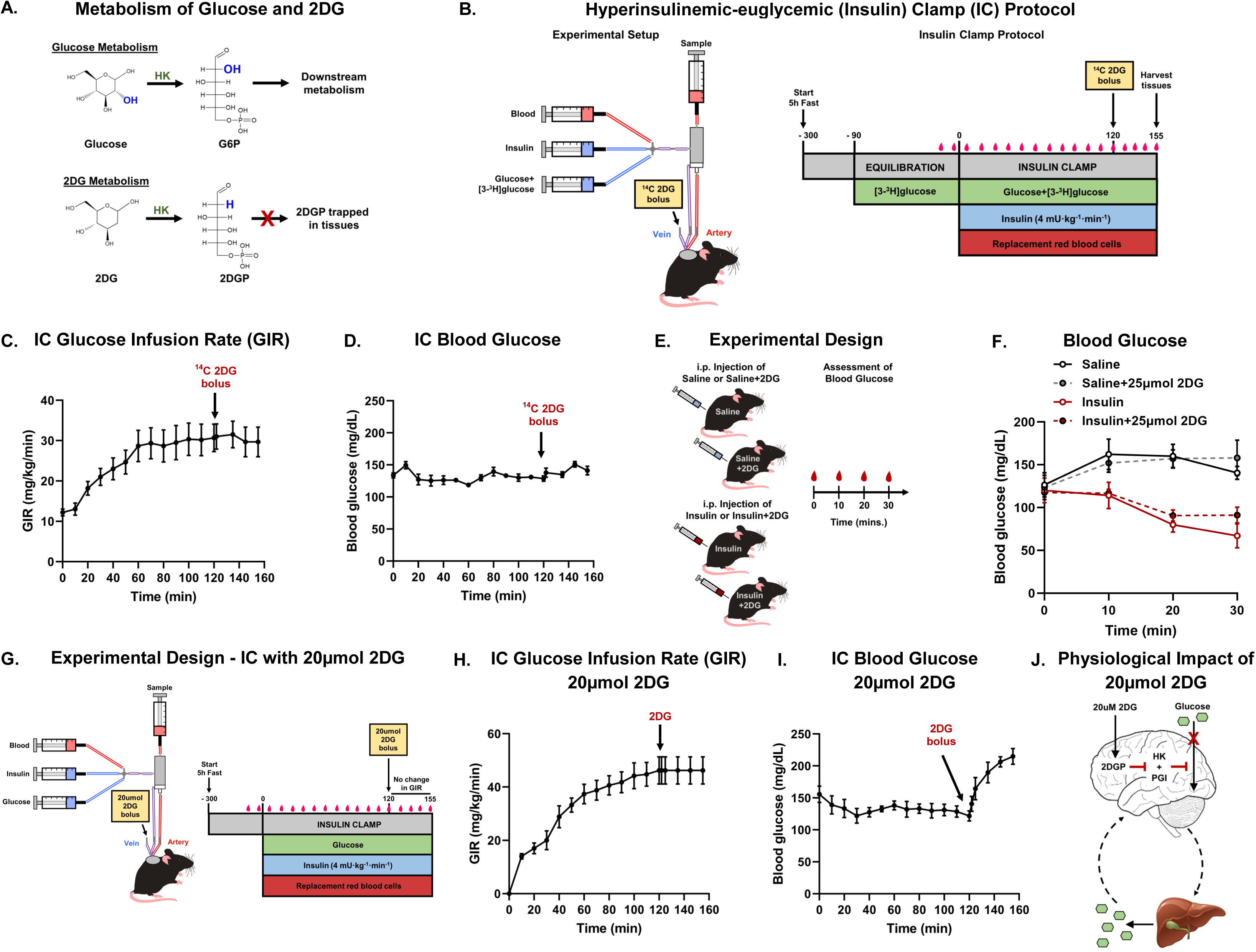
Measurement of *in vivo* glucose uptake requires nanomolar quantities of glucose analogs. (**A**) Metabolism of glucose and 2-deoxyglucose (2DG). (**B**) Standard hyperinsulinemic-euglycemic (insulin) clamp (IC) protocol with radiolabeled tracers in mice. (**C**) Glucose infusion rate (GIR) during the insulin clamp (IC) with radiolabeled tracers. (**D**) Blood glucose during the insulin clamp (IC) with radiolabeled tracers. (**E**) Experimental design for assessment of blood glucose in mice injected with saline or insulin with or without 25 µmol 2DG. (**F**) Blood glucose from mice injected with saline or insulin with or without 25 µmol 2DG. (**G**) Experimental design for assessment of blood glucose after venous administration of 20 µmol 2DG during an insulin clamp (IC). (**H**) Glucose infusion rate (GIR) during the insulin clamp (IC) with venous administration of 20 µmol 2DG. (**I**) Blood glucose during the insulin clamp (IC) with 20 µmol 2DG. (**J**) Schematic of the proposed physiological effect of venous administration of 20 µmol 2DG during an insulin clamp. (**C**-**D**, **F**, and **H**-**I**) Data are mean ± SEM. (**C**-**D**) N=6. (**F**) N=4 per group. (**H**-**I**) N=5. N represents biological replicates.

### 3.2. Validation of 2FDG as a suitable tracer for determining tissue-specific glucose uptake in mice

Our previous work demonstrated that unlabeled 2DG is poorly suited as an *in vivo* tracer for LC-MS-based assays due to interfering ions in biological samples, such as plasma and skeletal muscle tissues [14]. As a result, we recently developed a LC-MS method that can quantify trace amounts of the non-radiolabeled glucose analog, 2FDG, and its phosphorylated metabolite, 2FDGP in biological samples after injection of only 225 nmol 2FDG [14]. To validate that our LC-MS method was suitable for the determination of glucose uptake in mice, we performed a dose response experiment (Figure 2A). Accordingly, male C57BL6/NJ mice were injected with two different doses of 2FDG: 75 or 225 nmol. As anticipated, our results demonstrated that mice injected with 225 nmol 2FDG, when compared to mice injected with 75 nmol, exhibited an approximate 3-fold increase in circulating 2FDG and a similar increase in tissue 2FDGP (Figures 2B and 2C). Notably, the tissue distribution of 2FDGP levels shown in Figure 2C was similar to those obtained using radiolabeled tracers [3] and, importantly, tissue levels of 2FDGP after a 225 nmol injection were well within the 2FDGP standard curve (Supplemental Figure 1A). Next, we sought to determine if 225 nmol 2FDG altered blood glucose during an insulin clamp (Figure 2D). To this end, 2FDG was administered during an insulin clamp and the GIR remained unchanged similar to the experimental design described in Figure 1G and Results section 3.1 (Figures 2D and 2E). Results showed that venous injection of 225 nmol 2FDG did not alter whole-body glucose metabolism during an insulin clamp (Figures 2E and 2F). Taken together, these data demonstrate our 2FDG method could be used to determine which tissues differentially take up glucose relative to surrounding tissues when exposed to the same glycemic environment.

**Figure 2.**
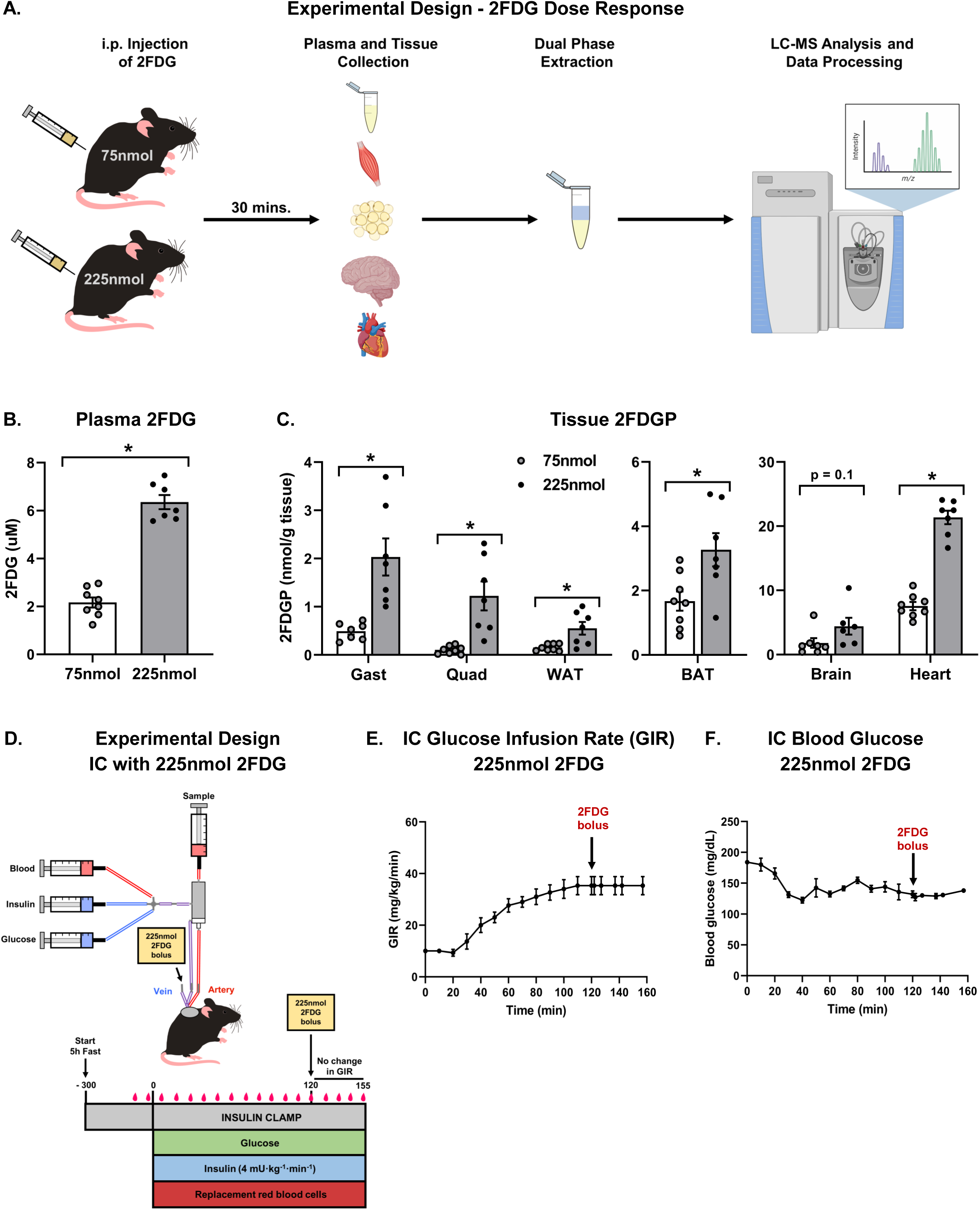
Validation of 2FDG as a suitable tracer for determining tissue-specific glucose uptake in mice. (**A**) Experimental design for 2FDG dose response experiment in mice. (**B**) Plasma 2FDG. (**C**) Tissue 2FDGP. (**D**) Experimental design for insulin clamp (IC) with venous administration of 225 nmol 2FDG. (**E**) Glucose infusion rate (GIR) during the insulin clamp (IC) with venous administration of 225 nmol 2FDG. (**F**) Blood glucose during the insulin clamp (IC) with 225 nmol 2FDG. (**B**-**C** and **E**-**F**) Data are mean ± SEM. (**B**-**C**) N=7-8 per group. (**E**-**F**) N=3. (**B**-**C**) Data were analyzed by two-tailed student’s t-test. (*) significant difference between 75 and 225 nmol. *P≤0.05. N represents biological replicates. See also Supplemental Figure 1.

### 3.3. Intraperitoneal injection of insulin with 2FDG is sufficient to detect differences in insulin-stimulated glucose clearance

One objective of this study was to develop a higher throughput, ‘catheter-free’ method to determine which tissues take up glucose in response to exogenous insulin. Among possible routes of administration, intraperitoneal (i.p.) injection is preferable to intravenous (i.v. or tail vein) injection because it provides reasonably rapid distribution into the circulation while requiring less specialized training. To determine whether i.p. injection was a suitable alternative for measuring insulin-stimulated glucose clearance, mice were injected i.p. or i.v. with 2FDG and either saline or insulin, followed by serial blood collection to measure blood glucose and plasma 2FDG. Tissues were harvested after 35 min (Figure 3A).

**Figure 3.**
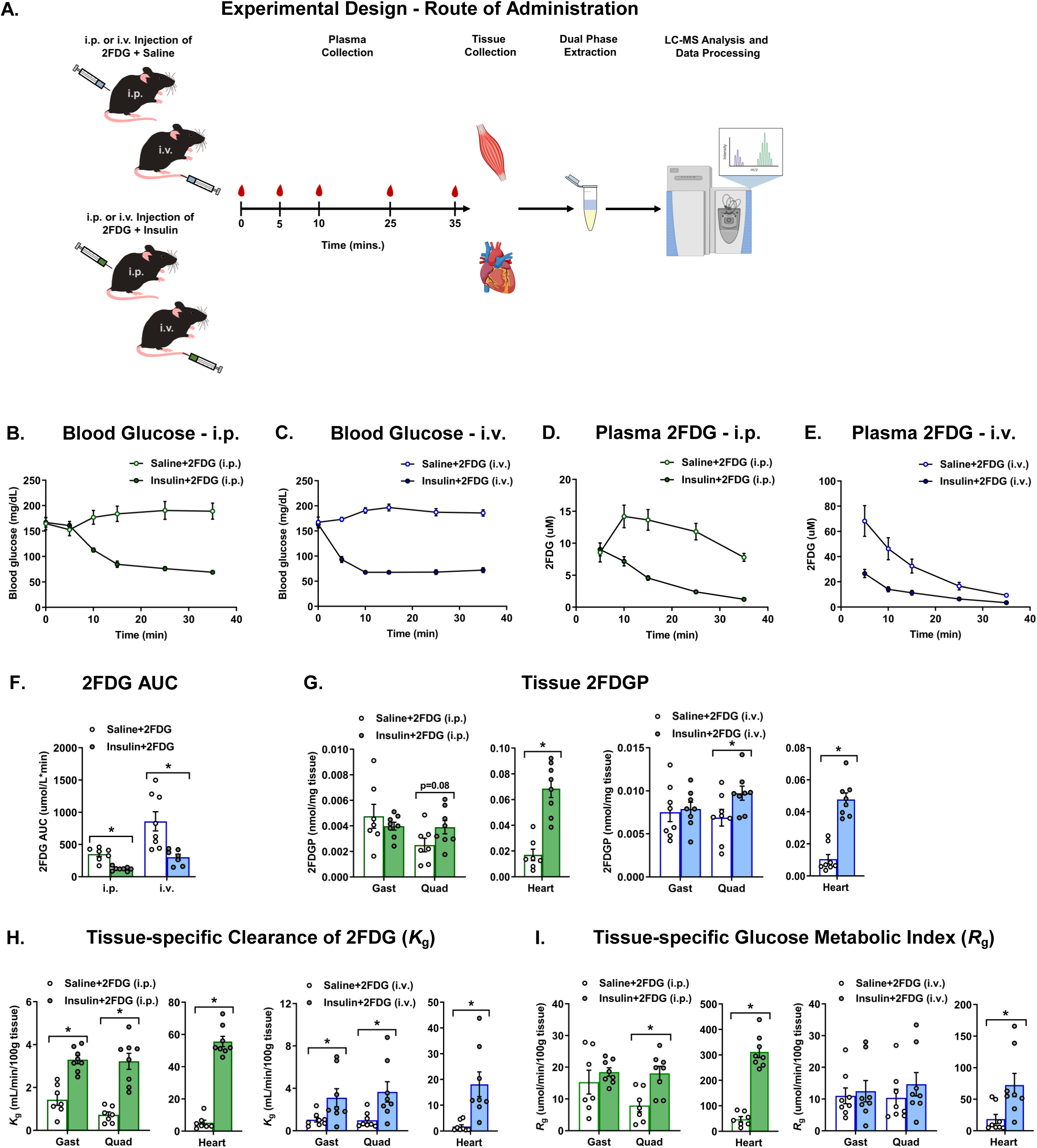
Intraperitoneal injection of insulin with 2FDG is sufficient to detect differences in tissue glucose clearance. (**A**) Experimental design for studies examining 2FDG route of administration. (**B**) Blood glucose after intraperitoneal (i.p.) injection of 225 nmol 2FDG with saline or insulin. (**C**) Blood glucose after intravenous (tail vein, i.v.) injection of 225 nmol 2FDG with saline or insulin. (**D**) Plasma 2FDG disappearance after intraperitoneal (i.p.) injection of 225 nmol 2FDG with saline or insulin. (**E**) Plasma 2FDG disappearance after intravenous (tail vein, i.v.) injection of 225 nmol 2FDG with saline or insulin. (**F**) Plasma 2FDG area under the curve (AUC). The green bars represent i.p. administration whereas the blue bars represent i.v. or tail vein administration. (**G**) Tissue 2FDGP. (**H**) Tissue-specific clearance of 2FDG (*K*_g_). (**I**) Tissue-specific glucose metabolic index (*R*_g_). (**B**-**I**) N=7-8 per group. Data are mean ± SEM and were analyzed by two-tailed student’s t-test. (*) significant difference between saline + 2FDG or insulin + 2FDG. *P≤0.05. N represents biological replicates. See also Supplemental Figure 2.

As expected, blood glucose and plasma 2FDG tracer kinetics depended on both the route of administration and whether insulin was present (Figures 3B–3F). When 2FDG and insulin were co-administered i.p., compared to i.v., the fall in blood glucose was delayed, and plasma 2FDG declined more slowly, consistent with slower entry of both 2FDG and insulin into the circulation via the i.p. route. To evaluate the effect of insulin on tissue glucose uptake, we focused on skeletal muscle (skm) and heart, tissues known to be major sites of insulin-stimulated glucose disposal. Regardless of how the tracer was delivered, insulin accelerated plasma 2FDG clearance, reflected by a lower 2FDG area under the curve (AUC) (Figure 3F). We next measured the accumulation of phosphorylated 2FDG (2FDGP), the trapped product of glucose uptake, in quadriceps (Quad), gastrocnemius (Gast), and heart (Figure 3G). Insulin increased 2FDGP in Quad and heart in both i.p.- and i.v.-injected mice; however, the effect in Quad was modest, and no difference in Gast 2FDGP was observed (Figure 3G). This result was unexpected, because skm is the primary site of insulin-stimulated glucose disposal [19].

Before interpreting this finding, it is important to recognize a fundamental limitation of tissue 2FDGP as a standalone measure: it reflects not only the rate of glucose uptake, but also how much tracer was available to be taken up and phosphorylated. If less tracer reaches the tissue, for example, because plasma 2FDG levels are low, then 2FDGP will be low regardless of how actively the tissue is transporting glucose. In other words, 2FDGP conflates the rate of uptake with tracer availability. Tracer availability is captured by the tracer/tracee ratio, or the ratio of 2FDG to glucose (2FDG/glucose) in plasma. When the tracer/tracee ratio is low, this indicates there was less tracer available for phosphorylation and thus measures of 2FDGP alone are not an accurate reflection of the rates of glucose uptake. To better understand why we did not observe the anticipated insulin-stimulated increase in skm 2FDGP, we next calculated the tracer to tracee ratio (2FDG/glucose) in plasma as a metric of the amount of tracer available for phosphorylation (Supplemental Figures 2A and 2C). When 2FDG was co-administered with insulin, the tracer/tracee ratio (2FDG/glucose) was lower than in saline-injected mice, especially in i.p.-injected animals, indicating that less tracer was available for phosphorylation. This means that tissue 2FDGP alone cannot accurately reflect the rate of glucose uptake when insulin is present and actively lowering blood glucose.

To begin correcting for tracer availability, tissue-specific glucose clearance (*K*_g_) is calculated by dividing tissue 2FDGP by the plasma 2FDG AUC. *K*_g_ normalizes tissue tracer accumulation to the total tracer exposure of the tissue, partially removing the confound of differing tracer availability between animals or conditions. Using this approach and combining MS-based measures of tissue 2FDGP with plasma 2FDG kinetics, we found that *K*_g_ was higher with insulin regardless of route of administration (Figure 3H). This confirms that insulin did stimulate glucose uptake in skm and heart, and that the earlier absence of a robust 2FDGP signal in Quad and Gast reflected insufficient tracer availability rather than a lack of insulin action. However, because *K*_g_ uses only the plasma 2FDG AUC, it does not account for how much unlabeled glucose is competing with the tracer at any given time, that is, it does not incorporate the tracer/tracee ratio (2FDG/glucose). This distinction matters particularly when insulin is present, because insulin simultaneously lowers glycemia and accelerates 2FDG clearance, causing the tracer/tracee ratio to shift dynamically over time. To account for these differences in absolute glycemia, the glucose metabolic index (*R*_g_) extends *K*_g_ by additionally adjusting for plasma glucose concentration, providing a more complete estimate of the absolute rate of glucose uptake (Figure 3I). However, when saline- and insulin-injected mice were compared, *R*_g_ and *K*_g_ gave discordant results (Figures 3H and 3I). To investigate this, we considered that both plasma glucose and the tracer/tracee ratio (2FDG/glucose) changed dynamically over time in insulin-injected mice, not just between groups at a single time point (Supplemental Figures 2A and 2C). Neither *K*_g_ nor conventional *R*_g_ fully captures this time-varying tracer/tracee ratio. We therefore calculated a modified index, *R*_g_’, by dividing tissue 2FDGP by the AUC of the 2FDG/glucose ratio, directly incorporating the time-varying tracer/tracee ratio into the denominator. When compared across conditions, *R*_g_’ agreed with tissue 2FDGP and with *R*_g_ for most studies, whereas *K*_g_ diverged from both, consistent with its failure to account for absolute glycemia and the changing tracer/tracee ratio (Supplemental Figures 2B and 2D). Nonetheless, *K*_g_ and *R*_g_ showed reasonably good agreement across both routes of tracer administration (i.p. vs. i.v.), supporting their use in guiding the design of subsequent experiments.

### 3.4. Validation of the 2FDG method for measurement of insulin-stimulated glucose uptake in lean and obese mice

Considering that high fat (HF) feeding induces skm insulin resistance characterized by decreased insulin-stimulated glucose clearance, we proceeded to determine if we could use our 2FDG assay to detect differences in insulin-stimulated glucose clearance between mice fed a standard chow diet or an obesogenic, HF diet (Figure 4A). Mice fed a HF diet were compared against a control group fed a standard chow diet. As expected, body mass was higher in HF fed mice (Figure 4B) and this was associated with more fat mass and slightly less lean mass compared to chow fed controls (Figure 4C). Next, we assessed markers of whole-body glucose homeostasis. Consistent with previous reports [20, 21], mice fed a HF diet exhibited higher fasting blood glucose (Figure 4D) and marked intolerance to an oral glucose challenge (Figures 4E and 4F). Since defects in skm glucose metabolism greatly contribute to insulin resistance, we hypothesized that impaired glucose tolerance in HF fed mice was due to decreased skm glucose clearance. To this end, we utilized our 2FDG method and assessed insulin-stimulated *K*_g_ and *R*_g_. After insulin injection, the rate of fall in blood glucose and plasma 2FDG levels were lower in HF fed mice (Figures 4G-4I) consistent with decreased clearance of glucose by peripheral tissues. As the data suggested differences in peripheral glucose clearance, we next sought to determine which tissues took up glucose in response to an insulin challenge. Accordingly, we measured 2FDGP in skm and heart and found that 2FDGP was lower in gastrocnemius (Gast) and quadriceps (Quad) muscles from HF fed mice compared to chow fed controls, whereas there was no difference in heart 2FDGP (Figure 4J). Next, we calculated skm and heart *K*_g_ and *R*_g_ (Figures 4K and 4L). As predicted, skm *K*_g_ and *R*_g_ were decreased in HF fed mice compared to chow fed controls. In contrast, despite differences in heart *K*_g_ between groups, there was no difference in heart *R*_g_. To better understand why *K*_g_ did not agree with *R*_g_ in the heart, we calculated the plasma 2FDG/glucose ratio and *R*_g_’ (Supplemental Figures 3A-3B). Results from these calculations showed that there was good agreement between 2FDGP, *R*_g_, and *R*_g_’ further reinforcing that *K*_g_ is influenced by differences in the glycemic environment.

**Figure 4.**
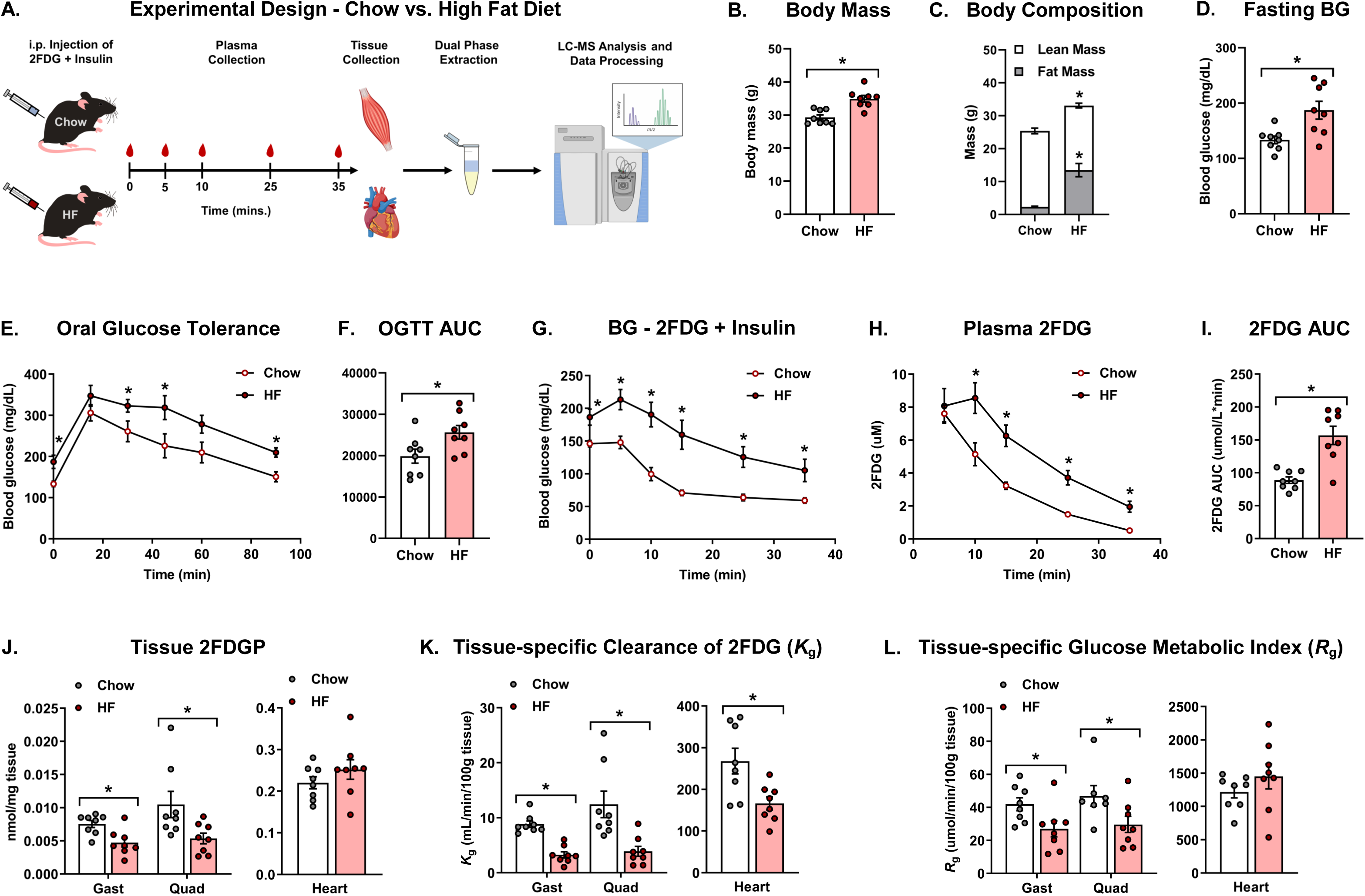
Validation of the 2FDG method for measurement of insulin-stimulated glucose uptake in lean and obese mice. (**A**) Experimental design for diet studies. (**B**) Body mass. (**C**) Body composition. (**D**) 5 h fasting blood glucose (BG). (**E**) Oral glucose tolerance. (**F**) Oral glucose tolerance test (OGTT) area under the curve (AUC) adjusted for baseline glucose. (**G**) Blood glucose after injection of 225 nmol 2FDG+insulin. (**H**) Plasma 2FDG disappearance. (**I**) Plasma 2FDG area under the curve (AUC). (**J**) Tissue 2FDGP. (**K**) Tissue-specific clearance of 2FDG (*K*_g_). (**L**) Tissue-specific glucose metabolic index (*R*_g_). (**B**-**L**) N=8 per group. Data are mean ± SEM and were analyzed by two-tailed student’s t-test. (*) significant difference between chow and high fat (HF) diet. *P≤0.05. N represents biological replicates. See also Supplemental Figure 3.

### 3.5. Application of the 2FDG method confirms increased insulin-stimulated skeletal muscle glucose uptake in TXNIP-deficient skeletal muscles

Mice lacking skm thioredoxin-interacting protein (TXNIP) are an established model of redox imbalance and enhanced skm insulin action ([22], Supplemental Figures 4A-4C). Considering the accepted role of TXNIP in the regulation of insulin-stimulated skm glucose metabolism, we proceeded to apply our 2FDG method to investigate insulin-stimulated tissue glucose clearance in chow fed mice lacking skm TXNIP (TXNIP^Myo^ or mKO) and littermate controls (TXNIP^fl/fl^ or FC) (Figure 5A and Supplemental Figure 4A). As predicted, there was no difference in body weight or body composition between the genotypes (Figure 5B and Supplemental Figure 4D). Next, we measured fasting blood glucose as a primary screen to evaluate if glucose homeostasis was different between genotypes and, as expected, our data showed that fasting blood glucose was decreased in mKO mice compared to controls (Figure 5C). Because of the foregoing difference in fasting blood glucose, we next assessed oral glucose tolerance. Herein, we show that there was a slight difference in the glucose excursion following oral administration of glucose (Figure 5D); however, when the glucose curves were corrected for differences in fasting glucose, the AUC revealed there was no significant difference in glucose tolerance between genotypes (Figure 5E). As differences in insulin during the OGTT and/or a liver phenotype could mask tissue-specific differences in glucose metabolism, we next deployed the 2FDG assay to assess insulin-stimulated glucose clearance. In the presence of insulin, there was no difference in blood glucose (Figure 5F); nevertheless, there was a slight, yet non-significant increase in plasma 2FDG clearance in mKO mice compared to controls (Figures 5G and 5H), suggestive of increased glucose clearance by peripheral tissues. Next, we applied the 2FDG method to quantify skm glucose clearance. As anticipated, mKO mice had a pronounced increase in insulin-stimulated Gast and Quad 2FDGP, *K*_g_, and *R*_g_ compared to controls (Figures 5I-5L). By contrast, 2FDGP, *K*_g_ and *R*_g_ were not different in the brain or heart between genotypes (Figures 5I-5L). These data support other studies showing that TXNIP specifically regulates muscle insulin action [23, 24] and, most importantly, further validate our method.

**Figure 5.**
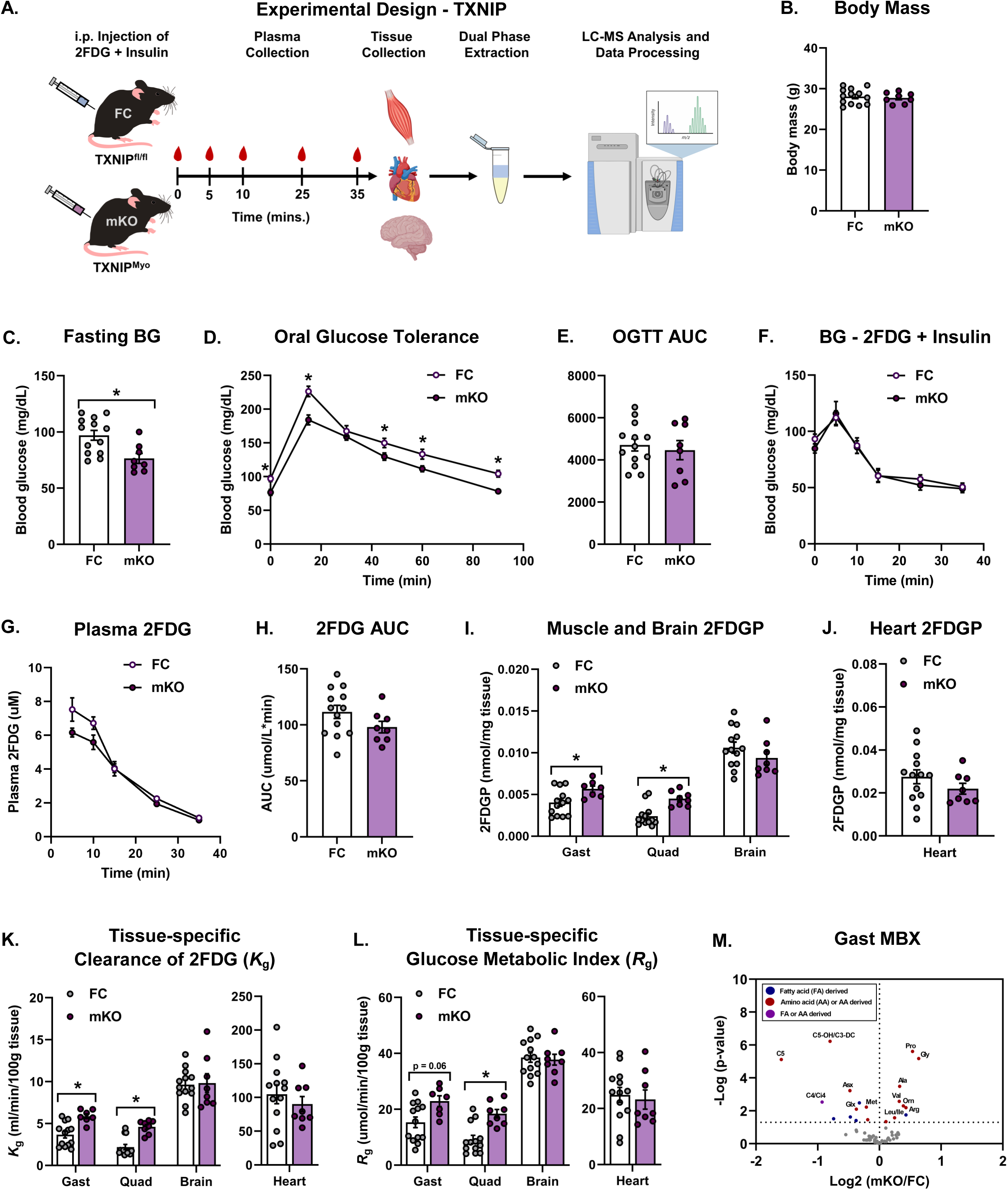
Application of the 2FDG method confirms increased insulin-stimulated skeletal muscle glucose uptake in TXNIP-deficient skeletal muscles. (A) Experimental design for studies in chow fed TXNIP^Myo^ (mKO) and floxed controls (FC). (B) Body mass. (C) Fasting blood glucose. (D) Oral glucose tolerance. (E) Oral glucose tolerance test (OGTT) area under the curve (AUC) adjusted for baseline glucose. (F) Blood glucose after injection of 225 nmol 2FDG + insulin. (G) Plasma 2FDG disappearance. (H) Plasma 2FDG area under the curve (AUC). (I) Muscle and brain 2FDGP. (J) Heart 2FDGP. (K) Tissue-specific clearance of 2FDG (*K*_g_). (L) Tissue-specific glucose metabolic index (*R*_g_). (M) Volcano plot of gastrocnemius (Gastroc) metabolites (MBX). (A-L) N=7-13 per group. (M) N=8 per group. Data are mean ± SEM. Data were analyzed by two-tailed student’s t-test. (*) significant difference between FC and mKO mice. *P≤0.05. N represents biological replicates. See also Supplemental Figure 4.

Considering the foregoing differences in insulin-stimulated skm glucose uptake, we next sought to determine the effect of TXNIP on the tissue metabolome to identify molecular signatures that best predict alterations in glucose homeostasis. To this point, targeted metabolomic profiling was performed in skm and liver using the exact same tissues collected from the 2FDG studies. Results showed that there was a shift in skm amino acids and amino acid-derived acylcarnitines in mKO mice compared to controls (Figure 5M and Supplemental Figure 4G). Several amino acids such as proline, glycine and the branch chain amino acids (BCAAs) - valine, leucine and isoleucine - were more abundant, whereas amino acid-derived acylcarnitines (AC) such as C5 and C5-OH/C3-DC AC were less abundant. Increased skm BCAAs is consistent with prior reports showing that skm TXNIP deficiency leads to decreased protein expression of the BCKAD E1α subunit and decreased BCKAD enzyme activity [22]. In contrast, there was little to no difference in the liver metabolome between genotypes (Supplemental Figure 4H). Taken together, these data validate the diagnostic capability of the 2FDG method and highlight its ability to be combined with secondary “omics” analyses.

## 4. DISCUSSION

Traditional methods to determine insulin sensitivity and glucose uptake in mice, such as the hyperinsulinemic-euglycemic (insulin) clamp, rely on the use of indwelling catheters and radiolabeled tracers. The numerous safety, administrative and logistical restrictions of radiolabeled tracers can be overcome using MS-based approaches that deploy non-radiolabeled glucose analogues for estimating metabolic fluxes *in vivo*. In theory, a non-surgical, non-radiolabeled approach would provide an attractive ‘intermediate’ phenotyping test that can be performed alongside glucose or insulin tolerance tests to assess tissue-specific sites of glucose clearance, and critically, specimens from the same animals could be used for secondary analyses, including state-of-the-art “omics” technologies. To address this methodological gap, we developed a higher throughput, radiolabel-free method for interrogating tissue-specific insulin-stimulated glucose metabolism in unrestrained, conscious mice. The assay can be integrated with other MS-based approaches to facilitate molecular characterization of the same tissue samples via discovery technologies such as genomics, proteomics and metabolomics. Beyond these analytical advantages, the practical throughput of the current method represents a meaningful improvement over the insulin clamp. A skilled investigator (with help) can typically complete insulin clamp experiments in approximately 4 mice per day, whereas the current method can be performed in approximately 8 mice per day. This throughput advantage makes the assay particularly well-suited as an intermediate phenotyping screen, one that can be performed alongside standard glucose and insulin tolerance tests to identify which tissues influence systemic glucose metabolism before committing to the more resource-intensive insulin clamp.

A key component of our assay is a modified, radiolabel-free version of the *in vivo* 2DG method. A significant consideration for the 2DG method is that high doses of 2DG (∼20 µmol) can inhibit hexokinase in the brain and significantly alter systemic blood glucose via a central counterregulatory response [18]. Whereas our results showed that an i.p. injection of 25 µmol unlabeled 2DG with or without insulin did not alter glucose metabolism, subsequent experiments using the insulin clamp revealed that a similar dose (20 µmol) significantly altered systemic blood glucose. Therefore, our data confirmed that nanomolar quantities of glucose analogs are required for the measurement of glucose clearance in rodents; and our findings underscore the importance of carefully evaluating the tracer load to ensure it does not perturb the metabolism under investigation.

As nanomolar quantities of tracer are required to ensure glucose metabolism is not disturbed, we subsequently developed and validated a new, highly sensitive MS-based method to determine tissue-specific glucose clearance in mice following an injection of trace amounts of 2FDG (225 nmol) with or without insulin. Results from these studies revealed that the 2FDG/glucose (tracer to tracee) ratio in plasma determines tissue 2FDG phosphorylation. For example, in the i.p. saline vs. insulin study, when the tracer was injected with insulin (as compared to saline) there was less tracer/tracee in plasma; therefore, we did not observe the anticipated insulin-stimulated increase in tissue 2FDGP; and this is exactly what one would expect even if the actual amount of glucose taken up is increased. Importantly, we also recognize that differences in the 2FDG/glucose ratio between mice can be overcome by normalizing muscle 2FDGP to brain 2FDGP [25]. Nonetheless, these data emphasize that measures of plasma 2FDG and the glycemic environment are critical for interpretation and, further underscores that any measure of tissue 2FDGP needs to be interpreted in light of the glucose (or precursor) pool.

To further validate the ability of the method to detect differences in insulin-stimulated glucose clearance, we applied it to two mouse models with anticipated changes in insulin sensitivity and peripheral glucose uptake - HF fed mice and mice lacking TXNIP in skm (mKO mice). In contrast to the striking differences in glucose uptake between chow and HF fed mice, the TXNIP mKO mouse model exhibited more subtle differences in systemic glucose metabolism, and we observed no differences in the fall in glucose after insulin administration between genotypes. This provided a unique opportunity to demonstrate the ability of the 2FDG method to detect differences in tissue glucose metabolism in the absence of overt systemic differences in glycemia after insulin administration. Indeed, when we compared insulin-stimulated glucose clearance in chow fed mKO mice and controls, glucose clearance was increased in skm, but not in hearts or brains, from mKO mice further validating our method.

When measures of glucose clearance were integrated with targeted LC-MS metabolomics, we identified shifts in amino acid and amino acid–derived acylcarnitine profiles associated with increased skeletal muscle glucose clearance in mKO mice. Notably, this included reductions in amino acid-derived acylcarnitine species, such as C5-DC, C5-OH/C3-DC, and C4/Ci4, suggesting altered flux through branched-chain amino acid (BCAA) and related metabolic pathways. This finding is consistent with established interactions between substrate utilization and insulin sensitivity in skeletal muscle. While the glucose-fatty acid cycle classically describes competition between lipid and glucose oxidation, BCAAs have also been implicated in impaired insulin signaling and glucose disposal when chronically elevated [26]. In the context of the TXNIP mKO model, the shift in the amino acid and acylcarnitine profiles, particularly reductions in amino acid-derived acylcarnitines, even in the setting of modest elevations in BCAAs, suggests decreased accumulation of incomplete oxidation products and improved mitochondrial handling of BCAA-derived carbon. Whether these metabolite shifts are a cause or consequence of improved glucose clearance in this model remains an open question, but they illustrate an important advantage of the non-radiolabeled approach: by preserving tissue biochemistry for downstream "omics" analyses, the same samples that yield glucose clearance data can simultaneously generate mechanistic hypotheses about the pathways driving or responding to changes in glucose metabolism.

In conclusion, we developed and validated a new non-surgical, non-radiolabeled mouse phenotyping test to assess *in vivo* tissue-specific glucose clearance. We envision this test as a screening ‘intermediate’ phenotyping test that can be employed alongside glucose and insulin tolerance tests to assess which tissues influence systemic glucose metabolism. With this knowledge, one could generate hypotheses that could be tested with more sophisticated measurements, such as the insulin clamp. When combined with other “omics” technologies, this method opens the door to innumerable mechanistic questions and experimental approaches to identify novel candidate mechanisms of tissue insulin resistance.

### Limitations

The current investigation examined the utility of combining 2FDG with an insulin injection to estimate glucose clearance by peripheral tissues. Importantly, we acknowledge that a major limitation of our method is that measurements of glucose clearance are made under non-steady state conditions. To this point, we strongly recommend that data are interpreted in light of group differences in glycemia and plasma tracer over the experimental time course. A modification of the protocols could include administration of a glucose load with the insulin and tracer to minimize the fall in absolute levels of glucose and eliminate possible confounding effects of hypoglycemia between groups in insulin sensitive models. If insulin action is part of the interpretation, a measure of plasma insulin is also needed. To this point, some may argue that adding 2FDG to a glucose tolerance test, as opposed to an injection of insulin, is a better option, as this is analogous to the flooding technique, wherein the 2FDG/glucose ratio may remain relatively constant; however, this would need to be tested. Lastly, another limitation of this study is that all methods were developed and validated exclusively in male mice; whether these findings generalize to female mice remains to be established, and future studies should include both sexes to assess potential sex-dependent differences.

## Author’s contributions

**Guofang Zhang:** Conceptualization, Methodology, Validation, Formal analysis, Investigation, Resources, and Writing – Review & Editing. **Dorothy H. Slentz:** Investigation. **Louise Lantier:** Investigation and Writing – Review & Editing. **Owen P. McGuinness:** Conceptualization, Resources, and Writing – Review & Editing. **Deborah M. Muoio**: Conceptualization, Resources, Supervision, and Writing – Review & Editing. **Ashley S. Williams**: Conceptualization, Methodology, Validation, Formal analysis, Investigation, Writing - original draft, Visualization, Supervision, and Funding acquisition.

## Acknowledgements

The authors would like to thank members of the Muoio Laboratory for their thoughtful feedback regarding these data and Richard Krentz and Alec Chaves for their technical assistance with the TXNIP studies. We also thank the faculty and staff at the Vanderbilt Mouse Metabolic Phenotyping Center (VMMPC) for their technical support with the insulin clamp studies. We would also like to thank the DMPI Metabolomics Core Lab (supported by Diabetes and Endocrine Research Center grant P30 DK124723) for the metabolomics data.

---

**Supplemental Figure 1.**
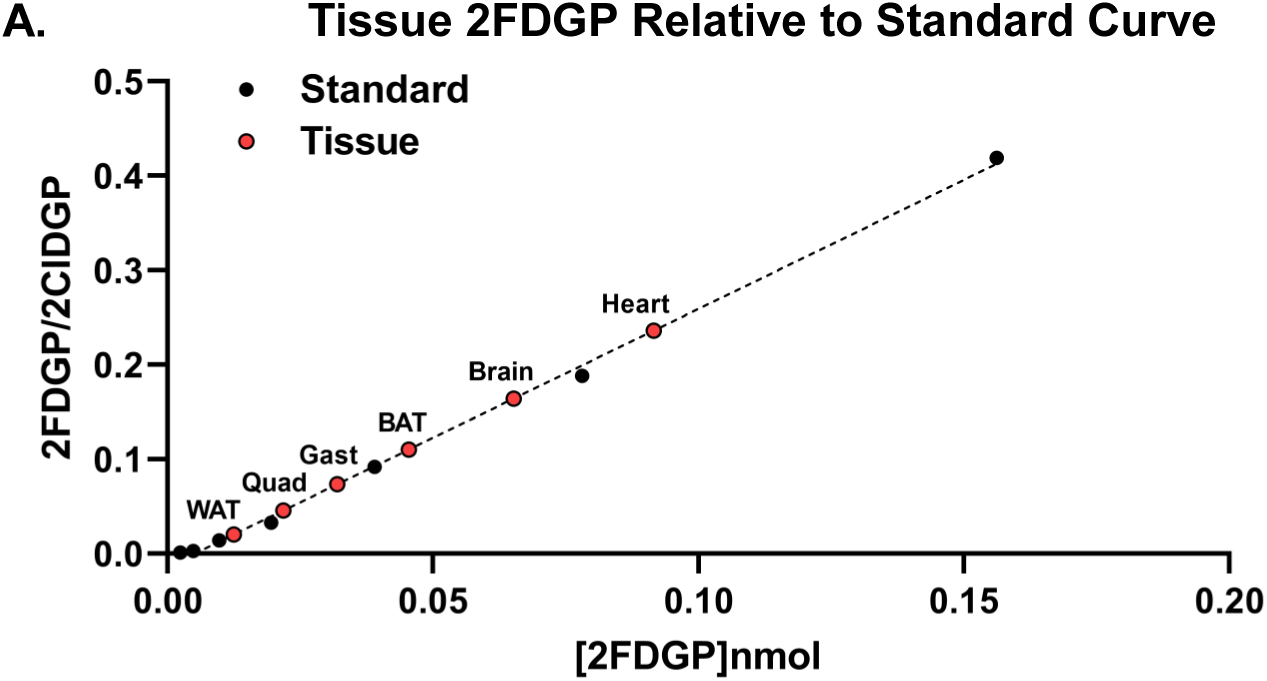
Tissue [2FDGP] relative to the 2FDGP standard curve. (**A**) Average tissue 2FDGP relative to the 2FDGP standard curve. Standards are in black and tissues are in red. Notably, the original 12-point standard curve was truncated to 7 points (0.156 nmol 2FDGP) to ensure all points were visible. Data are means of tissue samples (N=7/tissue). N represents biological replicates.

**Supplemental Figure 2.**
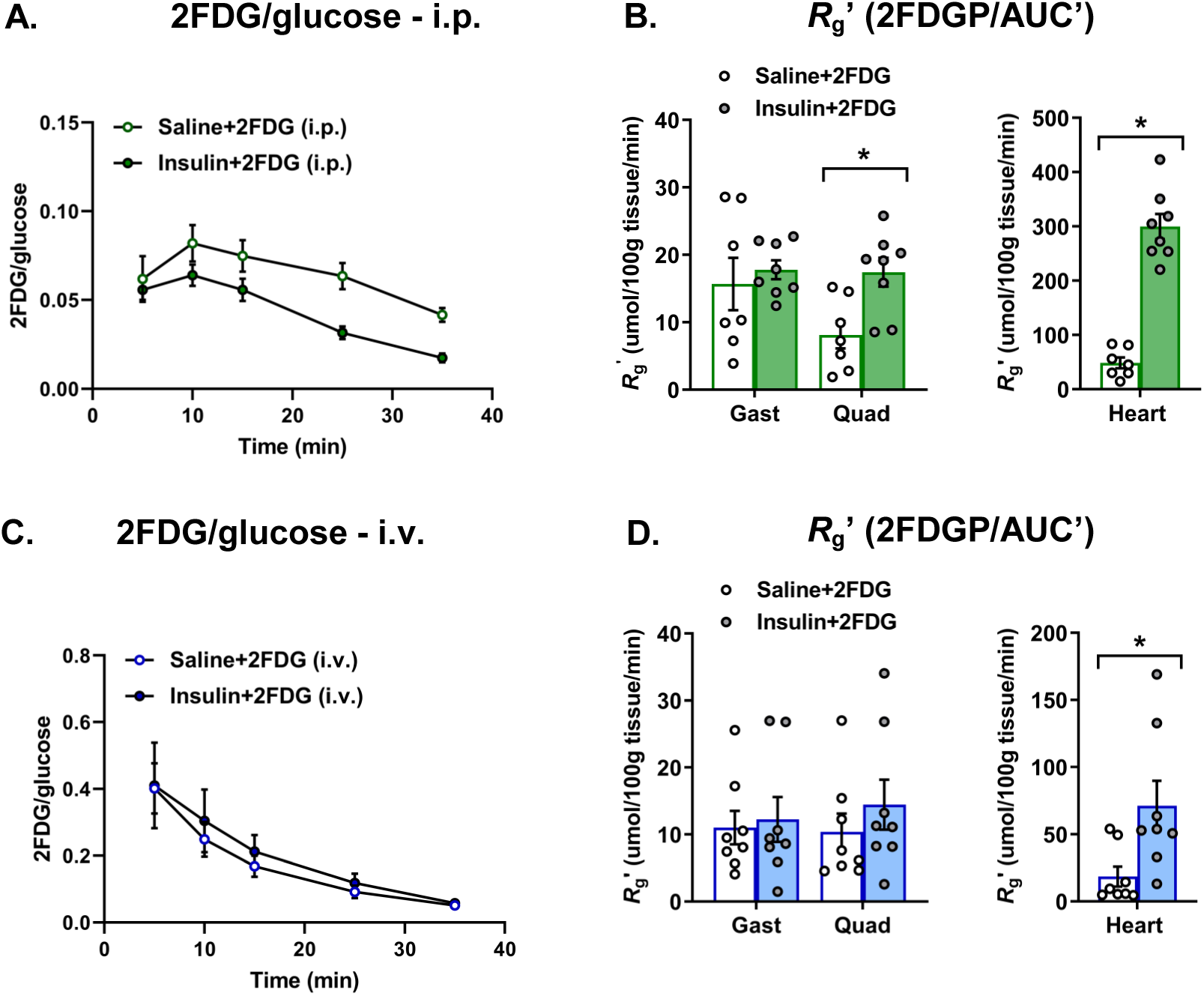
Plasma 2FDG/glucose and tissue *R*_g_’ in mice injected with 225nmol of 2FDG + saline or 225nmol of 2FDG + insulin. Ratio of plasma 2FDG to plasma glucose after (**A**) i.p. or (**C**) i.v. injection. Tissue *R*_g_’ (2FDGP/AUC’) after (**B**) i.p. or (**D**) i.v. injection. (**A**-**D**) N=7-8 per group. Data are mean ± SEM and were analyzed by two-tailed student’s t-test. (*) significant difference between saline + 2FDG or insulin + 2FDG. *P≤0.05. N represents biological replicates.

**Supplemental Figure 3.**
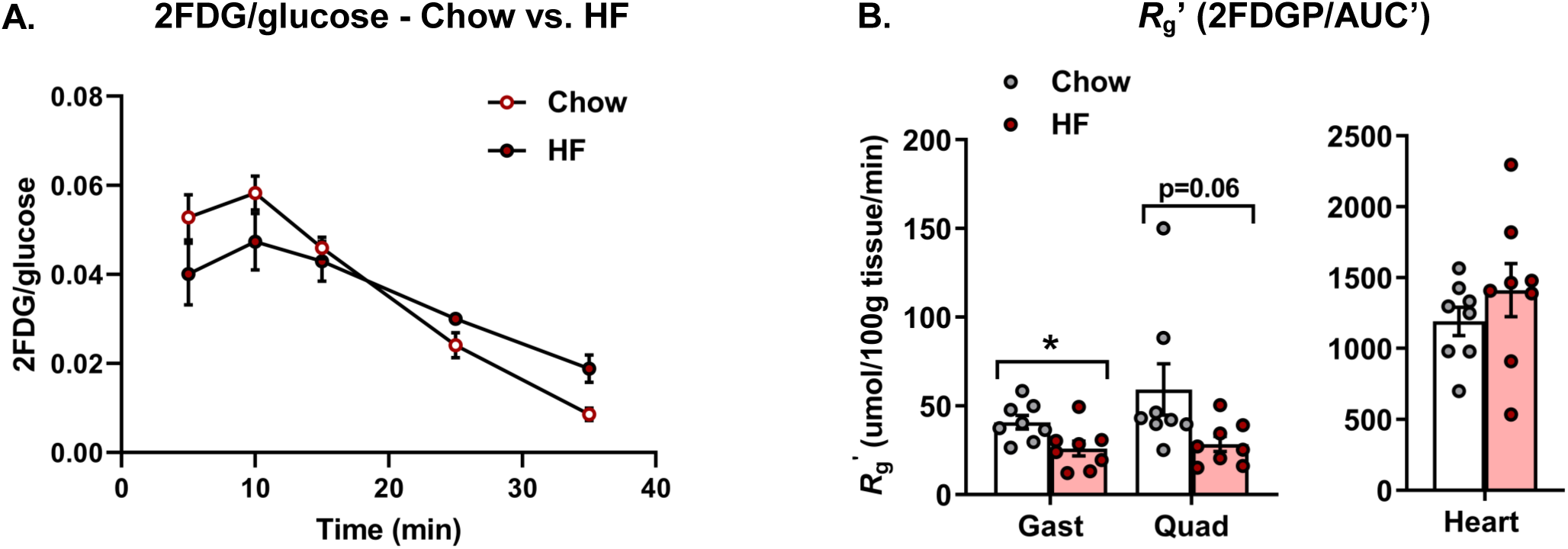
Plasma 2FDG/glucose and tissue *R*_g_’ in chow and HF fed mice injected with 225nmol of 2FDG + insulin. (**A**) Ratio of plasma 2FDG to plasma glucose and (**B**) *R*_g_’ (2FDGP/AUC’) after i.p. injection with 225nmol of 2FDG with insulin in chow and HF fed mice. (**A**-**B**) N=8 per group. Data are mean ± SEM and were analyzed by two-tailed student’s t-test. (*) significant difference between chow and high fat (HF) diet. *P≤0.05. N represents biological replicates.

**Supplemental Figure 4.**
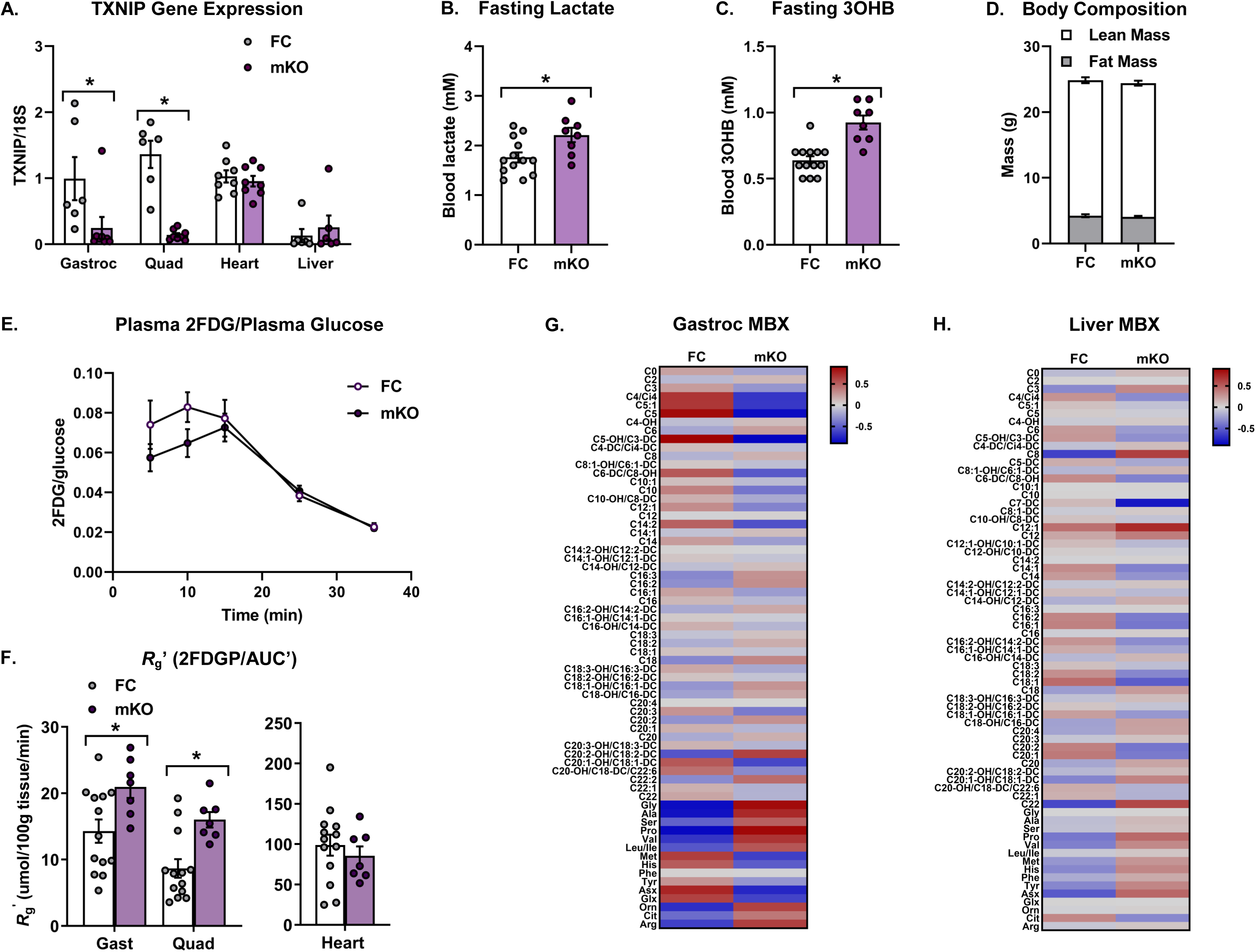
Phenotyping of chow fed TXNIP^Myo^ (mKO) mice. (**A**) TXNIP gene expression. (**B**) 5 h fasting lactate. (**C**) 5 h fasting beta-hydroxybutyrate (3OHB). (**D**) Body composition. (**E**) Ratio of plasma 2FDG to plasma glucose. (**F**) *R*_g_’ (2FDGP/AUC’). (**G**) Heatmap of gastrocnemius (Gastroc) metabolites (MBX). (**H**) Heatmap of liver metabolites. (**A**) N=6-8 per group. (**B**-**F**) N=7-13 per group. (**G**-**H**) N=8 per group. (**A**-**F**) Data are mean ± SEM. Data were analyzed by two-tailed student’s t-test. (*) significant difference between chow and high fat (HF) diet. *P≤0.05. (**G**-**H**) Data represent mean of the z-score. N represents biological replicates.

## REFERENCES

1. Ayala, J.E., et al., Considerations in the design of hyperinsulinemic-euglycemic clamps in the conscious mouse. Diabetes, 2006. 55(2): p. 390–7.

2. Ayala, J.E., et al., Standard operating procedures for describing and performing metabolic tests of glucose homeostasis in mice. Dis Model Mech, 2010. 3(9-10): p. 525–34.

3. James, D.E., K.M. Burleigh, and E.W. Kraegen, Time dependence of insulin action in muscle and adipose tissue in the rat in vivo. An increasing response in adipose tissue with time. Diabetes, 1985. 34(10): p. 1049–54.

4. Kraegen, E.W., et al., Dose-response curves for in vivo insulin sensitivity in individual tissues in rats. Am J Physiol, 1985. 248(3 Pt 1): p. E353–62.

5. Fueger, P.T., et al., Control of muscle glucose uptake: test of the rate-limiting step paradigm in conscious, unrestrained mice. J Physiol, 2005. 562(Pt 3): p. 925–35.

6. McGuinness, O.P., et al., NIH experiment in centralized mouse phenotyping: the Vanderbilt experience and recommendations for evaluating glucose homeostasis in the mouse. Am J Physiol Endocrinol Metab, 2009. 297(4): p. E849–55.

7. Williams, A.S., et al., Integrin alpha1-null mice exhibit improved fatty liver when fed a high fat diet despite severe hepatic insulin resistance. J Biol Chem, 2015. 290(10): p. 6546–57.

8. Kaihara, K.A., et al., PKA Enhances the Acute Insulin Response Leading to the Restoration of Glucose Control. Diabetes, 2015. 64(5): p. 1688–97.

9. Mottillo, E.P., et al., Lack of Adipocyte AMPK Exacerbates Insulin Resistance and Hepatic Steatosis through Brown and Beige Adipose Tissue Function. Cell Metab, 2016. 24(1): p. 118–29.

10. Cooney, G.J., et al., Insulin response in individual tissues of control and gold thioglucose-obese mice in vivo with [1-14C]2-deoxyglucose. Diabetes, 1987. 36(2): p. 152–8.

11. Wang, S.P., et al., Reply to Letter to the Editor: "The art of quantifying glucose metabolism". Am J Physiol Endocrinol Metab, 2017. 313(2): p. E259–E261.

12. Robciuc, M.R., et al., VEGFB/VEGFR1-Induced Expansion of Adipose Vasculature Counteracts Obesity and Related Metabolic Complications. Cell Metab, 2016. 23(4): p. 712–24.

13. Cutler, H.B., et al., Dual Tracer Test to Measure Tissue-Specific Insulin Action in Individual Mice Identifies In Vivo Insulin Resistance Without Fasting Hyperinsulinemia. Diabetes, 2024. 73(3): p. 359–373.

14. Williams, A., D. Muoio, and G. Zhang, A Fast and Sensitive Method Combining Reversed-Phase Chromatography with High Resolution Mass Spectrometry to Quantify 2-Fluoro-2-Deoxyglucose and Its Phosphorylated Metabolite for Determining Glucose Uptake. ChemRxiv, 2019. 2019(0227).

15. Kang, L., et al., Matrix metalloproteinase 9 opposes diet-induced muscle insulin resistance in mice. Diabetologia, 2014. 57(3): p. 603–13.

16. Varlamov, O., C.L. Bethea, and C.T. Roberts, Jr., Sex-specific differences in lipid and glucose metabolism. Front Endocrinol (Lausanne), 2014. 5: p. 241.

17. Wilson, R.J., et al., Disruption of STIM1-mediated Ca(2+) sensing and energy metabolism in adult skeletal muscle compromises exercise tolerance, proteostasis, and lean mass. Mol Metab, 2022. 57: p. 101429.

18. Borg, W.P., et al., Local ventromedial hypothalamus glucopenia triggers counterregulatory hormone release. Diabetes, 1995. 44(2): p. 180–4.

19. Thiebaud, D., et al., The effect of graded doses of insulin on total glucose uptake, glucose oxidation, and glucose storage in man. Diabetes, 1982. 31(11): p. 957–63.

20. Gnudi, L., et al., High level overexpression of glucose transporter-4 driven by an adipose-specific promoter is maintained in transgenic mice on a high fat diet, but does not prevent impaired glucose tolerance. Endocrinology, 1995. 136(3): p. 995–1002.

21. Turner, N., et al., Distinct patterns of tissue-specific lipid accumulation during the induction of insulin resistance in mice by high-fat feeding. Diabetologia, 2013. 56(7): p. 1638–48.

22. DeBalsi, K.L., et al., Targeted metabolomics connects thioredoxin-interacting protein (TXNIP) to mitochondrial fuel selection and regulation of specific oxidoreductase enzymes in skeletal muscle. J Biol Chem, 2014. 289(12): p. 8106–20.

23. Hui, S.T., et al., Txnip balances metabolic and growth signaling via PTEN disulfide reduction. Proc Natl Acad Sci U S A, 2008. 105(10): p. 3921–6.

24. Chutkow, W.A., et al., Deletion of the alpha-arrestin protein Txnip in mice promotes adiposity and adipogenesis while preserving insulin sensitivity. Diabetes, 2010. 59(6): p. 1424–34.

25. Halseth, A.E., D.P. Bracy, and D.H. Wasserman, Overexpression of hexokinase II increases insulinand exercise-stimulated muscle glucose uptake in vivo. Am J Physiol, 1999. 276(1): p. E70–7.

26. Newgard, C.B., et al., A branched-chain amino acid-related metabolic signature that differentiates obese and lean humans and contributes to insulin resistance. Cell Metab, 2009. 9(4): p. 311–26.

